# Three-dimensional patient-derived models of glioblastoma retain intra-tumoral heterogeneity

**DOI:** 10.1101/2025.01.08.632025

**Authors:** Zachery Moore, Claire Storey, Daniel V. Brown, Adam Valkovic, Montana Spiteri, Jasmine F. Pignatelli, Shannon J. Oliver, Alana Fakhri, Katharine J. Drummond, Seth Malinowski, Keith L. Ligon, Oliver M. Sieber, James R. Whittle, Saskia Freytag, Sarah A. Best

## Abstract

The intra- and inter-tumoral heterogeneity of glioblastoma represents a significant therapeutic challenge, as well as difficulty in generating reliable models for *in vitro* studies. Historical 2D adherent cell lines do not recapitulate this complexity, whereas both patient-derived neurospheres (PDN) and organoids (PDO) demonstrate intra-tumoral heterogeneity. Here, we quantify the tumor cell composition from matched models established from the same primary tumor using a series of multi-omic interrogations. We find that both patient-derived models recapitulate the genomic, epigenomic and tumor cell heterogeneity of the primary tissue. Furthermore, single-nuclei RNA sequencing revealed a subset of organoids containing small numbers of non-malignant cells from neuron and immune cell lineages. Harnessing the intra-tumoral heterogeneity of PDN models, we reveal the impact of temozolomide chemotherapy on individual cell states, altering composition of tumors over time in response to treatment. Our data confirms that both patient-derived models recapitulate patient intra-tumoral heterogeneity providing a platform for tumor cell state refined therapeutic studies.

**Key Points:**

- Generation of matched patient-derived neurosphere and organoid models from resected GBM tissue
- Neurosphere models exhibit greater proliferative signatures
- Both patient-derived models recapitulate genomic and epigenomic features of the primary tissue
- Single-nuclei RNA sequencing reveals both models recapitulate intra-tumoral heterogeneity of the primary tissue
- Neurosphere models enable interrogation of therapeutic responses in the context of heterogeneity

**Importance of study:** Patient derived models can be powerful tools when they faithfully recapitulate the tumor tissue from which they are derived. In glioblastoma, patient derived neurospheres (PDN) and organoids (PDO) have both been used in studies, however the differences between these models and the recapitulation of patient heterogeneity remain to be fully characterized. To address this, we performed multi-omic profiling of PDN and PDO models generated from the same tumor tissue. We find that that across a range of data modalities, both model systems exhibit a high level of resemblance to tissue, and critically, maintain heterogeneity and tumor cell composition. The importance of modeling heterogeneity was demonstrated in PDN models, where temozolomide treatment specifically alters the abundance of MES-like and AC-like tumor cells. Our findings demonstrate that neurosphere and organoid models effectively preserve cellular heterogeneity, genomic alterations, methylation signatures and transcriptomic features, both highly suitable to model glioblastoma’s complex cellular landscape.

## Introduction

The clinical management of glioblastoma (GBM) has remained unchanged since 2005, where the combined multimodal approach of resection, radiation and temozolomide chemotherapy was established^1^. Genomic characterization has revealed the mutational landscape of glioma^2^, with key alterations incorporated into updated WHO CNS classification^3^. However, this has not improved stratification of GBM into clinically relevant cohorts, with no targeted treatments demonstrating clinical benefit for GBM. Essential to the identification of effective treatments is the development of models that recapitulate the complex inter- and intra-tumoral heterogeneity observed in primary patient tissue. Failure of therapeutic translation using traditional cell line models of GBM can, in part, be attributed to these models that poorly account for such complexity. The use of clinically relevant, diverse *in vitro* models is crucial for therapeutic discovery and the elucidation of how individual tumors respond to therapies.

Intra-tumoral heterogeneity in GBM has been elucidated by the description of tumor cell states using transcriptomics, revealing close parallels to neurodevelopmental cell lineages^4,5^. Four tumor cell states are currently defined: astrocytic (AC-like), mesenchymal (MES-like), neuronal progenitor (NPC-like), and oligodendrocyte progenitor (OPC) lineages, with each displaying different biological behavior in GBM^4,6–8^. NPC-like and OPC-like cells have been shown to drive invasion through creation of neuron-glioma synapses^7^, whilst MES-like cells are involved in intricate cross-talk with the tumor immune microenvironment^9^. These tumor cell states are inherently plastic, with the states existing on a continuum and changing composition in response to treatment, as exemplified by an increase in MES-like population in response to temozolomide treatment^10,11^. Spatial and single-cell examination of individual tumors has revealed that despite common overrepresentation of an individual state, tumors are comprised of heterogeneous tumor cell states containing most, if not all, states.

Mapping of these states onto data from *in vitro* two-dimensional (2D) cell lines has revealed a poor recapitulation of these diverse tumor cell states using bulk RNA sequencing (RNAseq) datasets^12^. In particular, 2D cell lines often exhibit enrichment of the MES-like phenotype over extended culture periods^13^. In contrast, three-dimensional (3D) tumor models derived from patient tissue more closely mirror the original tumor^12^. These models include patient derived organoids (PDO)^14^, generated from culturing small pieces of tissue in a shaking incubator, and regarded as the gold standard in 3D cell culture due to their ability to mimic *in vivo* tissue architecture. Alternatively, spheroid 3D models can also be generated from the complete dissociation of tissue to single cells followed by culture in serum free media, termed patient derived neurospheres (PDN)^15,16^, or historically, glioma stem cell cultures (GSCs)^17^. While both 3D models show closer relatedness to patient tissue^14,18–24^, matched models established from the same initial tumor have not been established or compared, limiting informed use of these patient-derived model systems.

Here, we address this shortcoming through a detailed multi-omic comparison of matched PDN, PDO and their respective primary patient tissue, as the ground-truth. Our studies highlight principle differences between these two models, including the enhanced capacity of PDNs for expansion, and the partial maintenance of non-malignant cells in PDO models. At the genomic and epigenetic levels, both models maintain copy number alterations and DNA methylation patterns observed in the matched tumor tissue. Through detailed tumor cell annotation at the single cell level, we show all major tumor states observed in tissue are represented in both models. Our work highlights that despite certain differences, both PDN and PDO models exhibit striking similarities in their recapitulation of the original tumor they are representing. Importantly, both models capture patient intra-tumoral heterogeneity and we demonstrated tumor cell state-refined use of PDNs in therapeutic development pipelines.

## Materials and Methods

### Human subjects

De-identified glioblastoma WHO CNS grade 4 glioma samples with matching blood were obtained from consented patients at the Royal Melbourne Hospital Neurosurgery Brain and Spine Tumour Tissue Bank (Melbourne Health Ethics # 341 2020.214) and analyzed under a protocol approved by the Walter and Eliza Hall Institute Research Ethics Committee (HREC 21/21).

### Generation of PDOs and PDNs

Fresh tissue was obtained in the laboratory within one-hour post-surgery and equally divided for PDO and PDN model development. PDOs were generated and maintained as described^14^. In brief, PDO models were cultured in PDO media (50% DMEM F12 (Thermo Fisher Scientific #11320033), 50% Neurobasal (Thermo Fisher Scientific #21103049), 1× GlutaMAX (Thermo Fisher Scientific #35050061), 1× Non-Essential Amino Acids Solution (Thermo Fisher Scientific #11140050), 1× Penicillin-Streptomycin (Thermo Fisher Scientific #15070063), 1× N2 supplement (Thermo Fisher Scientific #17502048), 1× B27 supplement without vitamin A (Thermo Fisher Scientific #35050061), 1× 2-mercaptoethanol (Thermo Fisher Scientific #21985023), and 2.5 μg/ml insulin (Millipore Sigma # I2643-25MG)), and culture ware placed on an orbital shaker within a humidified incubator (37°C, 5% CO_2_). PDO models were passaged via macro-dissection into ∼1 mm^3^ pieces according to visual assessment. PDN models were generated through a combination of enzymatic (Miltenyi Biotec #130-095-942) and mechanical tissue (Miltenyi Biotec #130-093-235) dissociation prior to seeding in ultra-low attachment culture ware (Corning #CLS3471). All PDN models were expanded in neurosphere culture medium (STEMCELL Technologies #05751) supplemented with growth factors (20 ng/mL human recombinant EGF (STEMCELL Technologies #78006), 10 ng/mL human recombinant bFGF (STEMCELL Technologies #78003), 2 μg/mL heparin (STEMCELL Technologies #07980) within a humidified incubator (37°C, 5% CO_2_). When required, PDN models were dissociated in Accutase (STEMCELL Technologies #07920) for 5 minutes at 37 °C and seeded at a minimum of 100,000 cells per well within a 6-well plate. Routine mycoplasma PCR testing was performed using a reaction mixture containing 5.0 μL of 2X GoTaq Green Master Mix (Promega, M7122), 0.5 μL each of forward and reverse primers for mycoplasma^25^ (Forward: 5’-GGGAGCAAACAGGATTAGATACCCT-3’, Reverse: 5’-TGCACCATCTGTCACTCTGTTAACCTC-3’) and human *PTGER* control (Forward: 5’-TACCTGCAGCTGTACGCCAC-3’, Reverse: 5’-GCCAGGAGAATGAGGTGGTC-3’) targets at a final concentration of 0.25 μM, 2.0 μL of nuclease-free water, and 1 μL of cell lysate to achieve a total reaction volume of 10 μL. PCR was performed using the following thermal cycling conditions: initial denaturation at 95°C for 5 minutes (1 cycle), followed by 35 cycles of denaturation at 95°C for 30 seconds, annealing at 55-58°C for 30 seconds, and extension at 72°C for 30 seconds, with a final extension step at 72°C for 5 minutes, and a final hold at 12°C.

### Nucleic acid extraction

Tissue pieces were snap frozen in dry ice and stored at -80 °C until use. Tissue was homogenized via either mortar and pestle cooled with liquid nitrogen followed by QIAShredder (Qiagen #79656) or via TissueLyser (Qiagen). All tissue samples were processed using the AllPrep DNA/RNA mini kit (Qiagen #80204) as per manufacturer instructions. Germline DNA was obtained from blood samples extracted using the QIAamp DNA Mini Kit (Qiagen #56304) as per manufacturer instructions.

### SNP array analyses

DNA was processed using the Infinium Global Screening Array v3 (Illumina #20030770). Input amounts ranged from 50-200 ng DNA due to varied availability (call rates >99% across all probes). Data was processed at the service provider (Australian Genome Research Facility) to produce a single report per sample analyzed. B allele frequencies and log-r ratios were then extracted from each report and processed using ASCAT^26^ 3.1.3, with germline samples acting as a reference for absolute CNAs.

### Methylation array analyses

DNA was obtained as described and processed via the Infinium Methylation EPIC v2.0 BeadChip Array (Illumina #20087706). For all methylation experiments, 200 ng of DNA was processed per sample. Due to low tumor purity, GL0128 samples were excluded. Data was processed using a combination of minfi 1.5.0^27^, limma 3.60.2^28^ and SeSAME 1.22.2^29^ packages. All samples exhibiting a mean detection p value < 0.05 were included in the analysis. Data then underwent a series of pre-processing steps, including dye bias correction, p-value with out of band array hybridization, and noob background subtraction. Problematic probes, including those as cross-reactive^29^, were removed. For comparison across model type, design matrices included model type and line as terms. Differential probe and region analyses were performed using SeSAME using default parameters. Probes were mapped back to genes for downstream GSEA analyses.

### Immunohistochemistry

PDOs and PDNs were first removed from wells, spun down (5 minutes, 200g, 4°C), resuspended in 1mL PBS, and then spun down once more. Fixation was performed by gently resuspending cells in 1 mL 4% PFA in PBS for 45 minutes at 4°C with intermittent mixing. After washing with PBS and centrifugation, cells were embedded in pre-heated Histogel, allowed to solidify, and transferred to a histology cassette in 70% ethanol before sectioning. Immunohistochemistry was performed at room temperature. Following dewaxing, slides underwent peroxidase block (3% H_2_O_2_) for 10 minutes, wash (0.05% Tween 20 (Merck Millipore #P1379) in PBS), 5% goat serum (Merck Millipore #G9023) block in 0.05% Tween 20 (Merck Millipore # P1379) in PBS for 1 hour, primary antibody incubation (1:1000 rabbit anti-GFAP (Cell Signaling Technologies #80788)), 1:400 rabbit anti-Ki67 (Cell Signaling Technologies #12202), 1:1000 rat anti-SOX2 (Thermo Fisher Scientific #14-6811-82)) in blocking solution for 1 hour, secondary antibody incubation (1:300 biotin anti-rabbit (SUPP #123), 1:300 biotin anti-rat (SUPP #123) for 30 minutes, incubation with ABC Reagent (Vector Laboratories, #PK-7100) for 30 minutes, and a final incubation with DAB (Vector Laboratories, #SK-4105) until signal was observable by eye. Slides were then cover slipped via the Leica CV5030 automated coverslipper. Stains for hematoxylin and eosin (HCE) were performed via the Leica AutoStainer ST5010. Images were obtained via Olympus cellSens 2.3 software on an Olympus CX23 upright microscope.

### Single nuclei/cell and bulk RNA sequencing

All samples were processed using the Parse Biosciences Evercode WT Mini v2 kit. Matched tissue, PDN and PDO cohorts were extracted as nuclei. Single cell suspensions of PDO and PDN lines were counted and centrifuged at 200 g for 5 minutes at 4 °C. Pellets were resuspended in lysis buffer (250 mM sucrose, 25 mM KCl, 5 mM MgCl_2_, 10mM Tris pH 8.0, 1mM DTT, 0.2 U/mL RNasin (Promega #N2511), 0.005 U/mL DNase I (Thermo Fisher Scientific # EN0521), 0.1% Triton X-100, sterile H_2_O). Extraction of nuclei was confirmed using the SYTOX Green nuclear stain (Thermo Fisher Scientific # S7020), with ∼ 100% efficiency across all lines. Nuclei were input into the Parse Biosciences Evercode Fixation Kit as per manufacturer instructions. Tissue pieces were transferred into a chilled Dounce homogeniser containing 1.5 mL of lysis buffer (as above). A series of strokes with both A and B pestles was used to dissociate tissue, with debris monitored by hemocytometer. A per-sample decision was made to undergo wash steps (200 g for 5 minutes at 4 °C, resuspend in 0.01 % BSA, 1 mM DTT, 0.2 U/mL RNasin, sterile DPBS) as excess washes cause nuclei loss. Single cells isolated from the TMZ exposure experiment were dissociated using Accutase, then single cell suspension was input into the Parse Biosciences Evercode Fixation kit as per manufacturer’s instructions. All samples were stored at -80 °C for < 6 months. Fixed cohorts were processed using the Evercode WT Mini v2 kit as per manufacturers protocols, with 2 individual sub-libraries composed of 10,000 nuclei each. Both sub-libraries were sequenced targeting 20,000 reads per nuclei (as per manufacturers protocol). Resultant FASTQ files were processed using the Parse Biosciences provided split-pipe 1.2.1 software package to perform alignment to the hg38 genome, and subsequent unique molecular identifier (UMI) counts with sample barcode demultiplexing. Individual sub-libraries were processed separately before being merged. CellBender 0.3.2^30^ was also used to test for doublets in the dataset. Mitochondrial expression and number of features identified for each nucleus was calculated using Seurat 5.1.0^31^ and filtered on a global limit of > 250 features and < 10 % mitochondrial genes. All data underwent suggested Seurat processing steps as outlined within provided vignettes. TIRE-Seq was performed and preprocessed as described^32^.

### Identification of non-malignant cells

Non-malignant cells were identified prior to data integration using numbat 1.4.0^33^, given the close relation of GBM cellular states to normal CNS cell types. For each set of matched PDN, PDO and tissue samples, we first pooled nuclei across each to produce a phased genome. We then used the tissue segmented genome as obtained from SNP data as a prior. All other parameters were kept as default values. For each set of matched models, we visualised all nuclei using a UMAP and re-classified malignant cells that clustered with non-malignant cells. We then calculated a UMAP for all nuclei combined, and again performed the same reclassification step. This approach facilitated the identification of non-malignant cells present in low abundance within their respective samples.

### Integration of snRNASeq data

scVI 1.1.2^34^ was used to integrate snRNASeq data to overcome batch and sample effects. The dataset was filtered to 5,000 most highly variable genes, we also set each individual donor as the batch to integrated on within the model setup, in addition to parameters *n_hidden = 128*, *n_latent = 50, dispersion = “gene-batch”* and *early_stopping = TRUE*. All other parameters were kept as default.

### Nuclei annotation

Annotation was performed using the SingleR 2.6.0^35^ package with malignant cell reference dataset from Couturier, et al. ^4^, and non-malignant from Ruiz-Moreno, et al. ^36^. Raw counts were normalized via Seurat with default parameters for test and reference datasets, then input to *SingleR* function. Annotation was performed on a per-cell/nuclei basis with default parameters.

### Single nuclei/cell RNA analysis

For cell cycle scoring, we utilized Seurat 5.1.0 to assign inferred phases to each individual nuclei with provided G2M and S. These scores were also used to regress out the effect of cell cycle in comparisons within individual cellular states. For RRHO analysis, DEG lists were ranked via a -log_10_(p) × sign(fold change) method, and input into the RRHO2 1.0^37^ package with default parameters. For distance network analyses, measures of integration was quantified using the network graph within Minimum Distortion Embedding using the scanpy *pp.neighbours* function, with a Gaussian method to ensure distances are calculated for all nuclei to each other. LASSO regression was performed via the glmnet package. For each imaging readout of interest, segmented values were used as the response variable, and normalized count matrices for filtered for the top 10% most variable genes were used as the input matrix. Alpha was chosen as 0.95, otherwise all parameters were kept as defaults. All DEG analyses were performed via Seurat using the DESeq2 method of identification. For all comparisons, relevant nuclei populations underwent pseudo-bulk aggregation before downstream analyses. For comparisons across model type, design matrix included model as a term, whilst comparisons across cellular states included both model and state terms. A significance threshold of FDR adjusted p value < 0.05 was used across all comparisons. For enrichment analyses, curated gene sets obtained from both MSigDB^38^ and the Gene Ontology Resource^39^. The fgsea1.30.0^40,41^ package was used to perform gene-set enrichment analyses of curated MSigDB sets. DEGs were ranked on a -log_10_(p) × sign(fold change) method. For Gene Ontology, we employed an overrepresentation approach via the enrichR 3.2^42^ package.

### Live-cell imaging

Dissociated PDNs were seeded at 1,000 cells per well in 40 μL media in a 384-well ultra-low attachment plate. 10μL of 50μM TMZ in media (0.1% DMSO) was added to each well to achieve a final concentration of 10μM once individual spheres formed and exponential growth was observed. Using the IncuCyte platform, images were taken every 4 hours for 8 days. A custom cellpose model was used in conjunction with the tracking software TrackMate to segment spheres and calculate a series of morphological measurements. Data was imported into R with a custom script to calculate growth rate (GR) values over time. To calculate doubling time within the IncuCyte, a set of wells were seeded with 2,000 cells and a logistic growth model was fit to both 1,000 and 2,000 cell wells, with shared top, bottom and doubling time parameters.

### CellTiter Glo

Following two doubling events within the IncuCyte, each plate underwent a CellTiter-Glo assay as per manufacturer’s instructions. 50μL of CellTiter-Glo 2.0 was added into each well (1:1 ratio) and allowed to incubate on a rotating orbital shaker (100RPM) for 5 minutes at room temperature. Each well was transferred to a white-walled 384-well plate and luminescence was measured on a plate reader (Promega, GloMax) with an integration time of 0.3 seconds.

### Quantification and Statistical Analyses

Statistical differences were evaluated using differential expression or methods otherwise stated in figures. Graphs were produced with ggplot2. All analyses were performed within R 4.4.1

## Results

### Patient derived models reffect genomic and epigenetic features of patient tissue

To examine the recapitulation of key characteristics of primary GBM tumors in 3D patient derived models, we generated matched PDN and PDO models from four patients (**Figure 1A**). From each surgically resected tumor, tissue was assigned to the generation of PDNs, PDOs^14^ and snap frozen for original tumor analysis. Blood was collected at the time of surgery to provide reference germline DNA for genomic investigation. All four tumors were WHO CNS^3^ Grade 4 GBMs, confirmed by histopathology and next generation sequencing (NGS), and represented both primary and recurrence cases, and *MGMT* promoter methylated and unmethylated tumors (**Table 1**). PDO models exhibited shorter establishment times (first passage: 23.25 ± 9.74 days) as compared to PDN models (first passage: 155.75 ± 63.55 days), with passage defined as either complete dissociation for PDNs or mechanical dissociation for PDOs for propagation (**Figure 1B**). Despite the extended time for PDN generation, established PDNs subsequently maintained a stable growth rate over serial passages (**Figure 1B**). Previous studies have shown that a limitation of PDOs is the ability for expansion and subsequent throughput^43^. This was confirmed in our study, where the considerable input requirements for downstream experiments, exhausted all PDO lines at passage two (**Figure 1B**).

**Figure 1.**
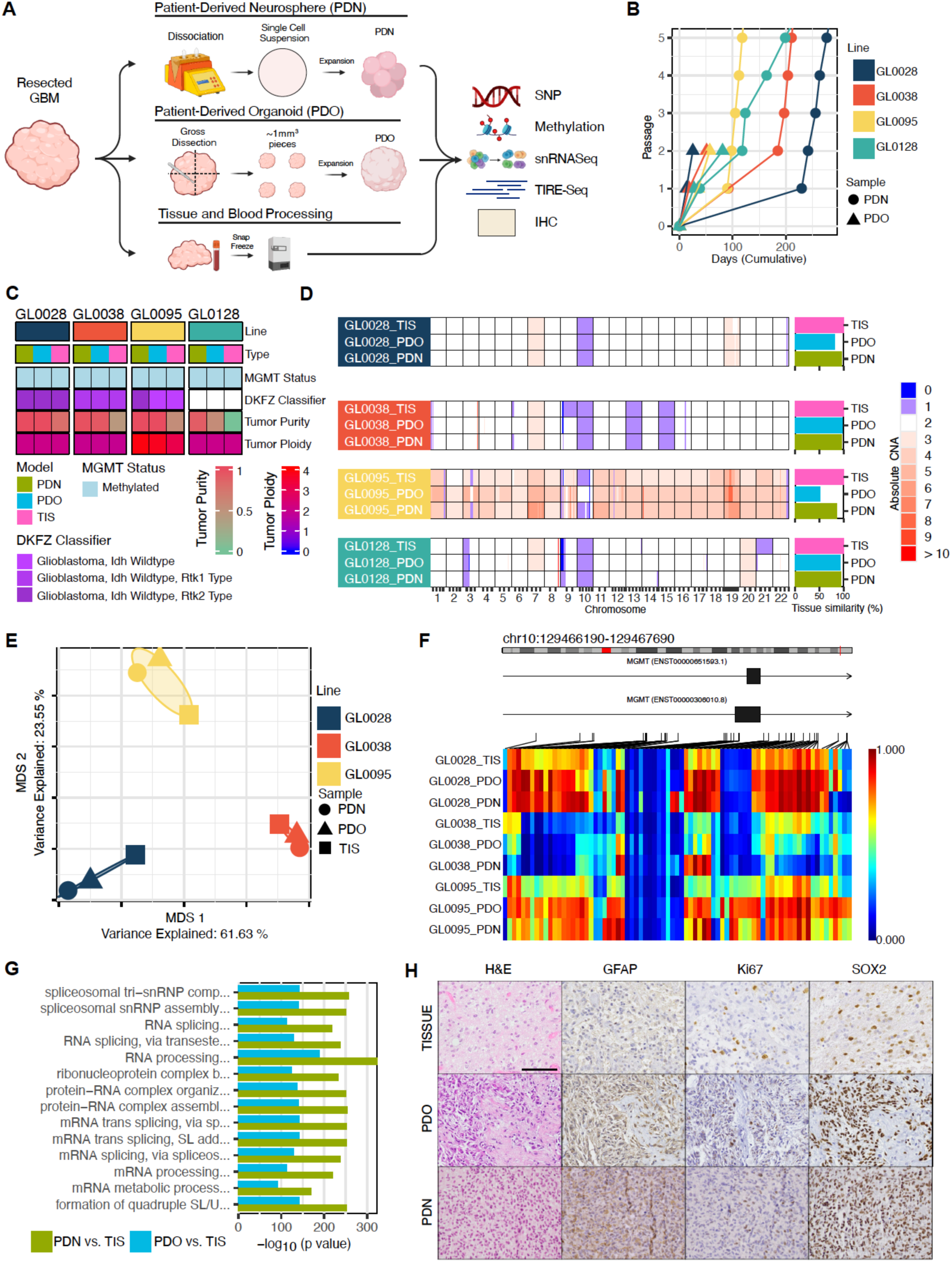
Genomic and epigenomic characterization of PDOs and PDNs shows broad concordance across models. **A.** Overview of study schema. **B.** Growth rates of individual models over time. A passage is defined as either a complete dissociation for PDNs or mechanical dissection for PDOs. Black circles indicate time when each model was processed for use in experimental investigations. Y-axis is limited to 5 passages, though PDNs do expand beyond this point. **C.** Heatmap of genomic and epigenomic characterization across each individual PDN, PDO, and tissue sample. **D.** Quantification of CNAs across individual lines and sample type as assessed via SNP array. Gains are shown red, losses in blue. All CNAs are absolute and compared back to a reference blood profile. Spearman correlation of CNAs across the entire genome within each individual line are shown on the right. **E.** Multi-dimensional scaling plot of EPICv2 methylation array data. Shapes represent individual sample types whilst colors represent individual lines. **F.** Visualization of β values obtained from EPICv2 arrays within the MGMT promoter region (1000bp before TSS to 500bp after), demonstrating concordance between tissue and PDN/PDO models within each individual line. **G.** Gene-set enrichment analysis of significant differentially methylated probes between either PDN and tissue or PDO and tissue. **H.** Representative IHC for GL0038 samples. IHC includes H&E stains, GFAP, KiC7, and SOX2 for PDN, PDO, and tissue samples. Scale bar, 100 μm.

**Table 1.** Clinical characteristics of individual samples and histopathological characteristics of tissue.

| **Clinical Characteristic** | **GL0028** | **GL0038** | **GL0095** | **GL0128** |
| --- | --- | --- | --- | --- |
| Diagnosis | GBM | GBM | GBM | GBM |
| Primary/Recurrent | Primary | Primary | Recurrent | Primary |
| Age | 42 | 62 | 51 | 38 |
| Sex | Male | Male | Female | Male |
| Extent of Resection | Gross macroscopic | Subtotal (50-95%) | Subtotal (50-95%) | Subtotal (50-95%) |
| Location | Occipital, temporal | Temporal | Frontal, parietal | Occipital, parietal, temporal |
| Previous Treatment | N/A | N/A | Stupp (3 rounds TMZ, 1 round 60 Gy RT) | N/A |
| *MGMT* promoter status | Methylated | Methylated | Methylated | Unmethylated |
| Ki67 quantification | ~ 20% | ~ 60% | Up to 30% | Up to 50% |
| GFAP | Positive | Positive | Positive | Positive |
| OLIG2 | Positive | Positive | Positive | Positive (Patchy) |
| p53 | Wild type | Wild type | Wild type (patchy) | Mutated |
| *TERT* promoter | Mutated | N/A | Mutated | Mutated |
| p16/CDKN2A | Lost | Lost | Lost | Lost |

To confirm the models reflected the genomic profile of the primary tissue, we investigated copy number alterations (CNA) in the tissue and patient derived models using single nucleotide polymorphism (SNP) arrays. Germline DNA extracted from the buffy coat of blood samples was used as a reference for each patient, enabling the detection of absolute CNAs and estimates of tumor purity. Initial examination of reference tumor tissue profiles revealed characteristic gains and losses of chromosomes 7 and 10, respectively, which were retained in both patient-derived models (**Figure 1D**). Further genome-wide examination of CNAs in PDNs and PDOs against tumor tissue profiles similarly revealed broad concordance in genomic alterations, although neither model was an exact replicate, as quantified via absolute correlation measures.

Of note, tumor GL0095 was found to be triploid, a rare event in GBMs which generally tend to be either diploid, or undergo a complete genome duplication to tetraploid^44^. This inherent genomic instability may explain why both GL0095 PDN and PDO models showed the lowest correlation measures against the original tumor for this case. Similarly, CNA profiles of PDN and PDO from GL0095 showed low correlation, suggesting outgrowth of genomically different subclones or model-specific divergence of each whilst within culture. Notably, initial analysis of GL0128 tumor tissue showed that this was largely comprised of non-malignant cells, with a calculated tumor purity of less than 5 % (**Sup. Figure 1A**). This hampered detection of CNAs in the tumor tissue, as compared to the models. Despite the low tumor purity, we nonetheless observed an *EGFR* amplification that was confirmed via NGS analysis and was also seen in the respective PDN and PDO models. Repeat SNP array analysis on a second tissue sample containing a higher proportion of tumor later revealed similar CNAs between the tumor and patient-derived models (**Figure 1D**). The low tumor purity from the original piece of tissue was reflected in both PDN and PDO models, where limited malignant starting material may have restricted the genetic diversity of overall tumor populations. Given GBM are heterogeneous, and all models are generated from distinct pieces of the resected tumor, it is encouraging to see both patient-derived models in agreement across the cohorts. Taken together, our data confirms that both PDNs and PDOs largely recapitulate genomic alterations found in original primary tumor tissue, despite models and reference tumor data having been generated from distinct pieces of the resected tumor.

Having confirmed that the models retained genomic features of the primary tumors, we next investigated if this held true for the epigenome. Global DNA methylation levels were measured using EPIC (version 2) arrays on primary tissue and 3D patient-derived models. Initial clustering showed clear separation of samples based on patient, suggesting that the 3D models all retained patient-derived methylation patterns in culture (**Figure 1E**). Methylation of the *MGMT*promoter region remains one of the few prognostic markers within GBM^1^. Examination of probes within this region (1,000 bp before and 500 bp after the transcription start site) revealed that all patient-derived models retained the respective *MGMT* methylation status of the patient tissue at the analyzed passages (PDO: passage 2; PDN: passage 3-4) (**Figure 1F**). We confirmed this result using an automated Methylation Classifier^45^. Furthermore, when using the classifier to predict the brain cancer subtype and grade, we found that the classification was consistent between all models and matched patient samples (**Figure 1C**), substantiating that both PDO and PDN models maintained the broad epigenetic signatures observed in the original patient tissue. Further investigation of the individual probes was performed using differential analyses to examine any differences between the patient-derived models and tissue. We identified only four significantly differentially methylated probes in matched PDNs compared to tissues, whilst no individual probe reached significance between PDOs and tissues. However, in both 3D patient-derived models several differentially methylated areas (DMAs) were detected compared to the tissue with greater divergence for PDN as compared to PDO models (10,034 DMAs in PDN vs. tissue, 2,177 in PDO vs. tissue, > 20 CpGs). We found that the bulk of DMAs covered the *HOX* gene family (**Sup. Figure 1B**), indicative of a more de-differentiated epigenome^46^. Gene set enrichment analyses (GSEA) performed for these DMAs identified 2,952 and 1,760 significant gene ontology (GO) terms for PDNs and PDOs respectively. Interestingly, the top 20 results for both models were related to increased levels of RNA transcription, with PDN comparisons showing more significant associations across sets (**Figure 1G**). This upregulation may be related to the accelerated cell proliferation of models in culture, which enhances metabolic and transcriptional activity, distinguishing these models from the slower-growing tissue environment. The more pronounced DNA methylation differences in PDN samples compared to PDOs may in part be due to longer passage time and higher passage number for PDN samples with increasing growth rate and adaption to culture media.

We then performed immunohistochemical (IHC) analyses to examine if these findings were reflected at the expression level (**Figure 1H**). Both models exhibited strong expression of glial fibrillary acidic protein (GFAP), and SOX2, a transcription factor associated with stemness and tumor-initiating potential. We also observed that Ki67, a proliferation marker, showed an increased proportion of positive cells in both PDN and PDO models. The spatial distribution and intensity of these markers in PDN and PDO models closely resembled those observed in primary tumor tissues. These similarities underscore the capacity of PDN and PDO models to capture the cellular and molecular heterogeneity of glioblastoma, making them valuable platforms for studying tumor biology and evaluating potential therapeutic strategies.

### PDO models can contain non-malignant cells

Next, to investigate the transcriptome across the cohorts and examine the tumor cell composition in patient-derived models compared to primary tissue, we performed single-nuclei RNA sequencing (snRNAseq). Following quality control steps, we obtained a total of 22,700 nuclei with a mean 3,123.16 genes expressed per nucleus. We first differentiated malignant and non-malignant cells interpolating inferred CNA using the numbat package^33^ to determine the composition of cells that grow in 3D culture environments. Given we had defined CNAs across the genome (**Figure 1C**), we used these as references to improve classification accuracy. We identified 2,882 non-malignant cells across the combined cohorts (**Figure 2A**). Consistent with previous genomic results, we confirmed that the initial GL0128 tissue sample was largely non-malignant, with our snRNAseq data composed of only 0.74% malignant cells. This skewed composition was reflected in the models generated from this mostly normal tissue, with the GL0128-PDO model an even mixture of non-malignant and malignant cells, and the GL0128-PDN model comprising a small number of non-malignant cells (**Figure 2B**). The presence of non-malignant cells in GL0128-PDN is an exception based on the input, whereas the three other models containing a more even mixture of tumor cells were composed entirely of malignant cells (**Figure 2B**). Interestingly, while PDO models are generally considered to all contain non-malignant cells, two models did not (GL0028 and GL0038). Given the low absolute numbers of malignant cells across all GL0128 samples, we excluded GL0128 models from subsequent analyses.

**Figure 2.**
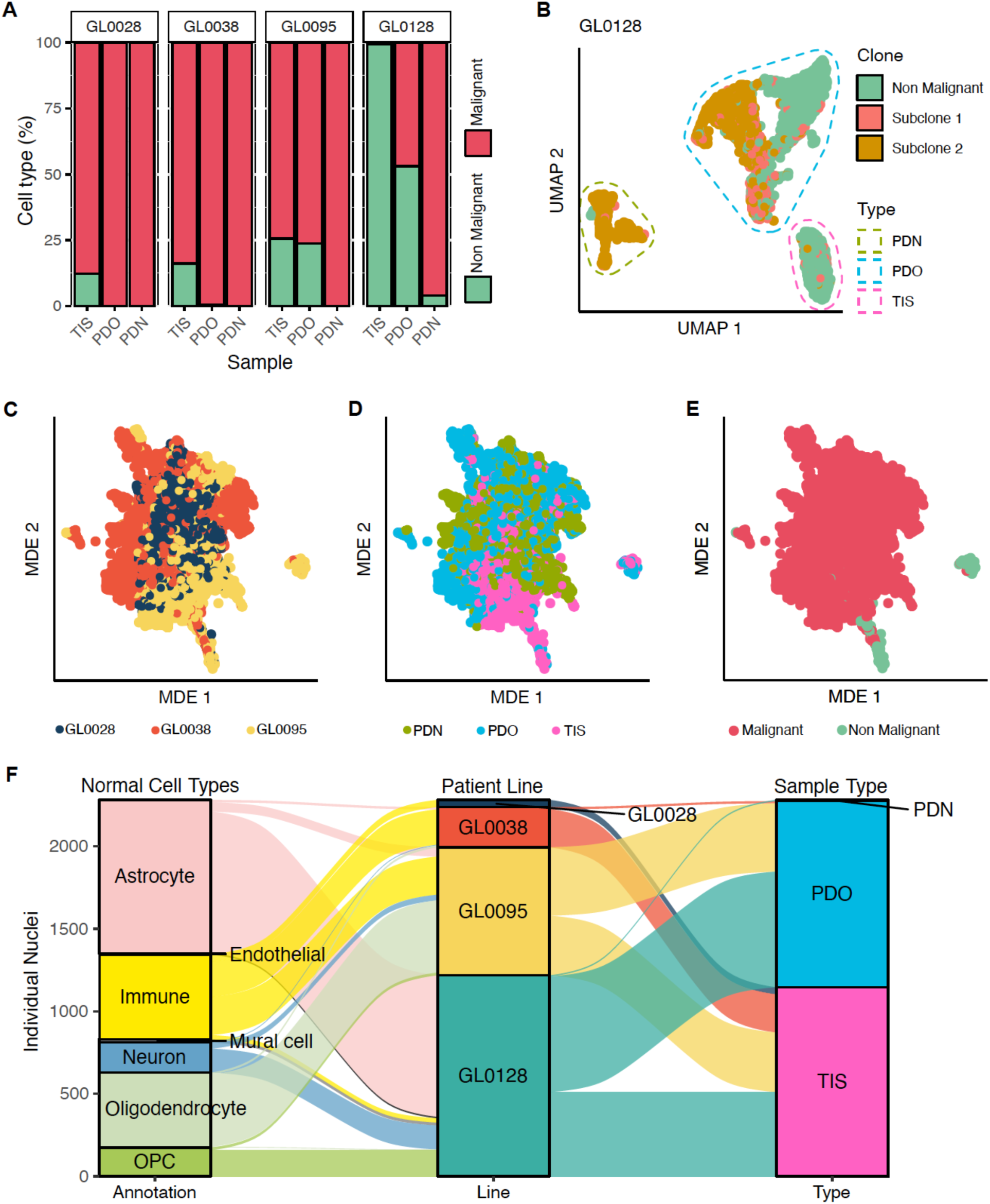
snRNASeq reveals both PDOs can contain non-malignant cells. **A.** Barplots of results obtained from numbat in identification of non-malignant cells separated by sample type and line. **B.** UMAP plot of GL0128 samples, demonstrating the predominant non-malignant composition of tumour tissue samples. **C.** Multi-dimensional embedding (MDE) plot of all snRNASeq data after integration with scVI demonstrating effective and homogenous integration across all samples. Each point represents an individual nucleus, with color representing line. **D.** As for (C), with color representing model type. **E.** As for (C), with color representing malignant and non-malignant cells as identified via numbat. **F.** Sankey diagram of re-annotated non-malignant cells reveals, first separated by non-malignant cell type, then individual line, and finally model type.

To further classify the non-malignant cells, we generated a unified space for downstream interrogation of data across all samples using scVI^34,47^ (**Figure 2C**). Cell annotation was confirmed using canonical marker gene expression within this integrated space, which validated the identity of the cell populations, including oligodendrocytes (*MBP*) and immune cells (*PTPRC*) (**Supp. Figure 2A, 2B**). All non-malignant cells were annotated based on reference-based methods using the dataset produced in *Ruiz-Moreno, et al.* ^36^. This revealed further non-malignant cell types within our dataset that included largely astrocytes, oligodendrocytes and immune cells, in addition to neurons, OPC cells and endothelium, which were exclusively derived from either PDO or tissue samples in each cohort (**Figure 2F**).

We find that in addition to GL0128-PDO, which we have demonstrated to be derived from a non-malignant piece of tissue, that GL0095-PDO, a recurrent GBM, also contained a high percentage (25%) of non-malignant cells. This may be in part due to the diffuse and invasive phenotype of recurrent GBM^48^, where malignant cells are interspersed within normal tissue. This finding further highlights the importance of tissue selection and the importance of carefully distinguishing tumor from normal cells within PDO cultures to ensure accurate characterization for downstream applications.

### PDNs exhibit upregulated proliferation signatures

To investigate the tumor cell composition and intra-tumoral heterogeneity in our patient-derived models, we next focused on the malignant cell populations. Pseudo-bulk analysis was first performed to investigate broad expression patterns between the primary tissue and patient-derived models. Overall, increased numbers of DEGs were identified between tissue and PDN (n = 252) compared to tissue and PDO (n = 111) (**Figure 3A, B**). We then compared the resulting DEG lists using rank-rank hypergeometric overlap (RRHO), finding a significant overlap (**Figure 3C**). This result indicates that overall, the difference between PDNs and tissue and PDOs and tissue are concordant, but more pronounced in PDNs. We then examined enriched GO sets that were upregulated in both models compared to the primary tissue. We found no significant enrichment for any terms within PDOs, but for PDNs observed significant enrichment for terms related to mRNA transcription and metabolism, in line with GSEA results for DNA methylation (**Figure 1G**). Given PDNs showed shorter times between passages, we then sought to determine if the observed increases in these signatures were due to increased tumor cell proliferation, focusing on progenitor populations which account for the majority of dividing cells in tumors, to measure the strength of this signal (**Figure 3D**). Consistent with this hypothesis, progenitor cells of both PDN and PDO models displayed significant GSEA enrichment for “Hallmark G2M Checkpoint Genes” compared to tissue (adjusted p = 3.8 × 10^−25^ and p = 1.1 × 10^−13^, respectively), with increased enrichment score (NES = 2.67 vs. 2.42) in PDNs compared to PDOs. Expression of *MKI67*, a canonical marker of cycling cells, was increased in all PDN models compared to PDO, which was also higher compared to tissue (**Figure 3E**). This result mirrors our findings at the protein level, where increased Ki67 abundance was observed in the PDN models (**Figure 1H**). We then used Cyclone to infer the cell cycle phase for all nuclei^49^. We observed that PDNs, PDOs and tissue all showed similar proportions of cells in each inferred cell cycle phase across tumor cell states (**Figure 3F**). This confirmed that the progenitor-like state contained the highest proportion of dividing cells amongst tumor states^4^. Thus, gene expression differences as compared to tumor tissue, including cell cycling signatures, are more pronounced in PDNs than PDOs, at least partly reflecting their increased capacity for proliferation.

**Figure 3.**
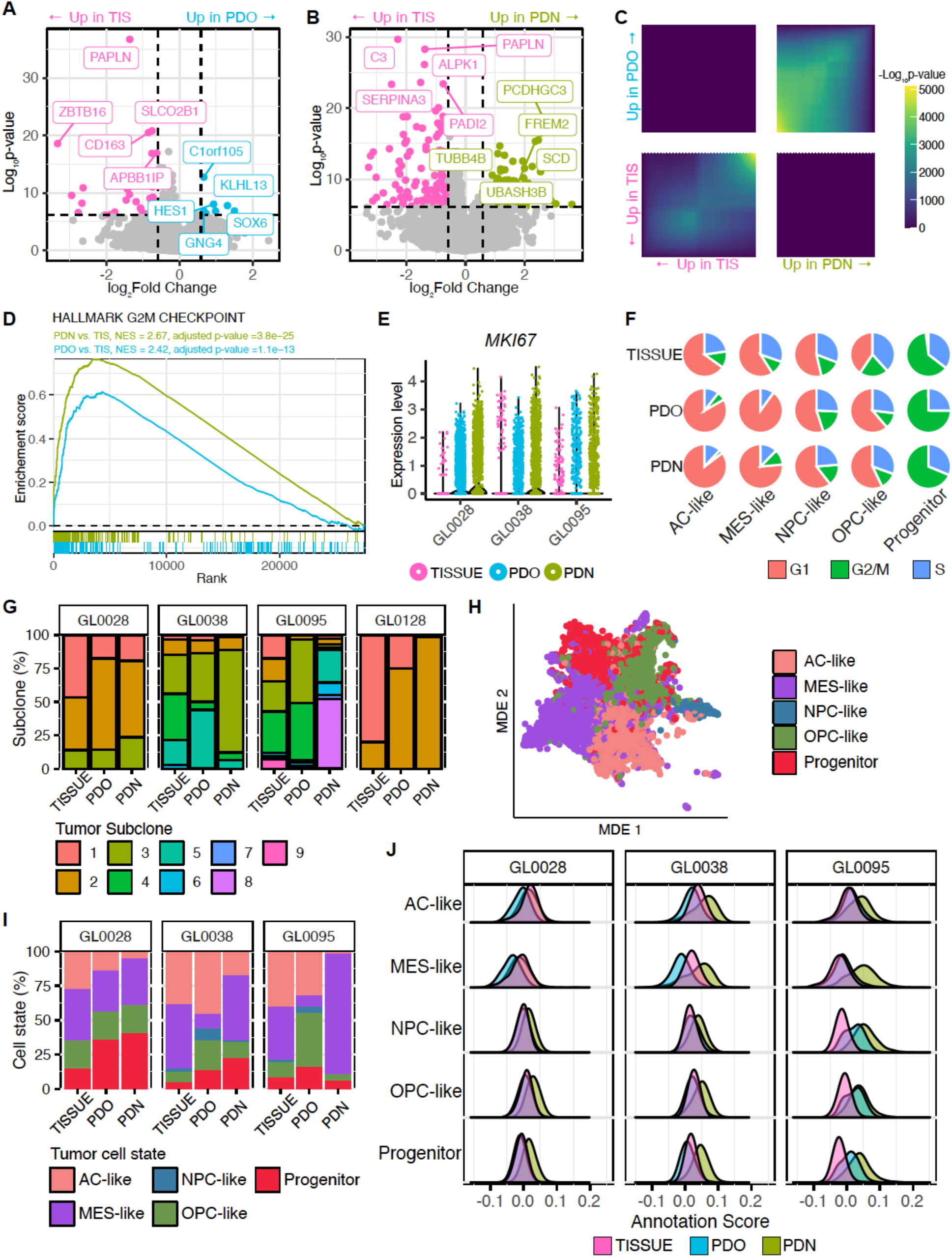
Tumor cell states are recapitulated in both PDO and PDN models. **A.** Volcano plots of DEGs between both pseudo-bulked malignant PDN nuclei compared back to malignant tissue nuclei. **B.** As for (A) but comparing malignant PDO nuclei compared to malignant tissue nuclei. **C.** Rank-rank hypergeometric plot of DEG lists obtained (A) and (B), showing overlap in gene ranks. Ranks determined by -log_10_(p_val) × sign(log_2_fold change). **D.** GSEA of HALLMARM_G2M_CHECKPOINT between PDN vs. tissue and PDO vs. tissue comparisons shown in (A) and (B) respectively. Ranks determined by -log_10_(p_val) × sign(log_2_fold change). **E.** Normalized expression levels of MKIC7 across all malignant nuclei, stratified by model type. **F.** Inferred cell cycle phase on snRNASeq data, separated on sample type and cellular state. **G.** Barplots of inferred subclonal compositions, separated by line and model type. Tumor subclones are not shared across lines and are exclusively composed of malignant cells. **H.** MDE plot of re-integrated tumour nuclei after scVI integration. Color represents individual cellular annotation. **I.** Barplot of best-assigned cellular annotation, separated by both line and model type. **J.** Distribution plots of annotation scores, separated by both line and sample.

### Subclonal heterogeneity is maintained in PDO and PDN models

Investigating sub clonal compositions across all individual lines (**Figure 3G**), found that most subclones detected in tissue were also detected in their respective matched models. GL0028 tumor tissue was found to be composed of three subclones, which were proportionally concordant across PDN and PDO models. In the GL0038 cohort, of six individual subclones found in tumor tissue, five were captured in PDN and PDO models. The inherent genomic instability of GL0095 was further reflected in the clonal investigation, with nine subclones detected across all models. Within this cohort, the PDN model was the most discordant compared to the tissue. The GL0128 tumor tissue contained only two subclones, illustrating a selective bottleneck in model generation from the low proportion of malignant cell input. While both subclones were identified in the patient-derived 3D models, their distribution was biased towards one subclone in both models, with the PDNs almost showing complete loss of the other subclone.

### Tumor cell states are recapitulated in both PDO and PDN models

To investigate tumor cell composition, we annotated our dataset based on the reference dataset produced in *Couturier, et al.* ^4^. We employed a semi-automated approach in our annotation pipeline, initially through a computational approach via the SingleR^35^ package, after which annotations were refined via manual inspection. This included expression of canonical markers, such as *TOP2A*, *CLU*, and *VIM* for progenitor, AC-like, and MES-like populations, respectively. We then integrated malignant cells using scVI in an unbiased manner, and confirmed our annotation as evidenced through clear separation of tumor states within the MDE space (**Figure 3H**). With tumor cell populations established, we next investigated the recapitulation of the patient-derived models to the ground-truth primary tumor sample within each cohort. Similar to tumor subclone proportions, broad concordance was observed between patient-derived 3D models and primary tissue with regards to proportions of tumor cell states (**Figure 3I**). All major states observed in tissue (>1%) were represented within each model. We also observed that broadly, both PDN and PDO models have enriched progenitor populations, which likely reflect their enhanced replicative ability compared to the primary tumor. A notable exception to concordant proportions was identified in GL0095 PDNs, which displayed an enriched MES-like population compared to both the PDO and primary tissue, suggesting a PDN-specific deviation. We also confirmed these results by calculating scores for tumor states originally defined in *Neftel, et al.* ^5^, and again showed concordance between models (**Sup. Figure 3**). Given tumor states display plasticity^4^ and our pipeline assigns tumor cell states using a best-fit method, we examined the distribution of assignment scores to further interrogate the differences between individual models (**Figure 3J**). We observed no significant difference between either PDN or PDO models in recapitulation of individual tumor states. As a surrogate measure for integration, we also extracted the distance metrics from the multi-dimensional nearest-neighbor network used in the creation of the MDE latent space. We find no significant differences across comparisons within a mixed-effects model, indicating equal integration between all sample types (**Sup. Figure 2D**).

We then expanded these analyses to interrogate DEGs within each individual tumor state between our patient-derived models. Once again, within each tumor state we observed a higher number of DEGs for PDNs versus tissue compared to PDOs versus tissue (AC-like: 167 vs. 50; MES-like: 195 vs.144; NPC-like: 0 vs. 2, OPC-like: 62 vs. 28; Progenitor: 120 vs. 28). However, a comprehensive KEGG pathway analysis to identify potential altered biological pathways found minimal differences across individual states, with six pathways for both comparisons reaching significance. These were seen across the MES-like and Progenitor populations for PDNs, and for AC-like, MES-like, and Progenitor populations for PDOs. PDNs displayed increased metabolism related pathways, whilst PDOs increased WNT and Notch signaling (**Supp. Figure 2E**). Taken together, this data suggests that the strongest signals that differentiates PDNs from PDOs are related to proliferation metabolism, reflected at both the bulk and individual tumor-state level.

### TMZ differentially impacts tumor cell states

Since PDNs recapitulated genomic, epigenomic and transcriptomic modalities and cell composition of the primary tumor with a greater capacity for expansion, we examined the response of tumor heterogeneity to therapy in PDN models. We conducted a combination of live-cell imaging and sequencing to understand the impact of temozolomide (TMZ) chemotherapy on tumor cell composition. We first performed a series of live-cell imaging experiments across six PDN lines, including the GL0095 model and five previously established PDN models (**Table 2, Supp. Figure 4A**). Each line was exposed to a range of TMZ concentrations from 100 nM to 100 μM. We observed a broad spectrum of responses to TMZ treatment, with the 50% inhibitory growth rate (GR_50_) ranging from 5.58 μM to 93.56 μM TMZ (**Table 2**). Notably, the GR_50_ and IC_50_ values exhibited no discernible correlation with cell doubling time (**Supp. Figure 4C**), indicating that the variability in drug sensitivity is not directly linked to proliferation rate. Using an independent GBM-derived PDN line (GL0073), we first treated at a single dose (100 μM TMZ or vehicle control) and analyzed heterogeneity at a single timepoint (**Figure 4A**). Single cell RNA sequencing (scRNAseq) was performed on the subsequent tumor cells to determine the impact of TMZ on tumor cell state heterogeneity and identify genes and pathways associated with resistance and sensitivity. UMAPs generated from the dataset revealed TMZ-treated cells intermixed with vehicle cells in all tumor cell populations (**Figure 4B**). Exposure to TMZ resulted in proportional increases to the MES-like tumor cells as previously reported^50^, though the largest increase was identified in AC-like tumor cells (**Figure 4C**). Examination of DEGs in the AC-like and MES-like populations between vehicle and TMZ treated cells revealed a common increase in the expression of splicing factors (*RNU4-2, RNU4-1)* (**Figure 4D**) and a negative enrichment for proliferation-related gene sets (**Figure 4E**) in TMZ-treated cells. To further investigate the longitudinal perturbation of tumor cell states as a result of TMZ, we utilized turbocapture integrated RNA expression sequencing (TIRE-seq), a high-throughput and low input method of RNAseq^32^ in parallel with live-cell imaging on the GBM BT504^16^ PDN line. This method enabled us to test a range of doses from 100 nM to 100 μM (7 doses and vehicle control) at three timepoints, 3-, 5- and 7-days post treatment (**Figure 4F**), aligning with 1, 1.5 and 2 doublings of the model. Initial PCA analyses showed clear separation of samples across both day and concentration (**Supp. Figure 4C**). Examining the impact of dose and time on tumor composition allowed us to determine a transcriptomic shift from 3.16 μM TMZ 5 days following treatment (**Figure 4G**). Paired live cell imaging data enabled us to use LASSO regression to identify genes most strongly associated with physical measures, including growth (area) and cell death (SYTOX and Caspase 3/7) (**Figure 4H**). *VEGFA,* a growth factor involved in blood vessel formation and cellular proliferation, demonstrated a positive correlation with PDN area from 5 days exposure. Increased concentrations of TMZ, which reduced PDN area, resulted in decreased *VEGF* expression. Cell death, measured by SYTOX green, was significantly correlated with *MMP2*, a matrix metalloprotease associated with poor outcomes in glioma^37^. *CHAC2* encodes a cyclotransferase protein involved in the degradation of glutathione, and whose decreased expression has been observed in several cancers^51^. We observe that increased *CHAC2* expression is seen in response to increasing TMZ concentrations, that decreases in expression are correlated with increases in Caspase 3/7 levels.

**Figure 4.**
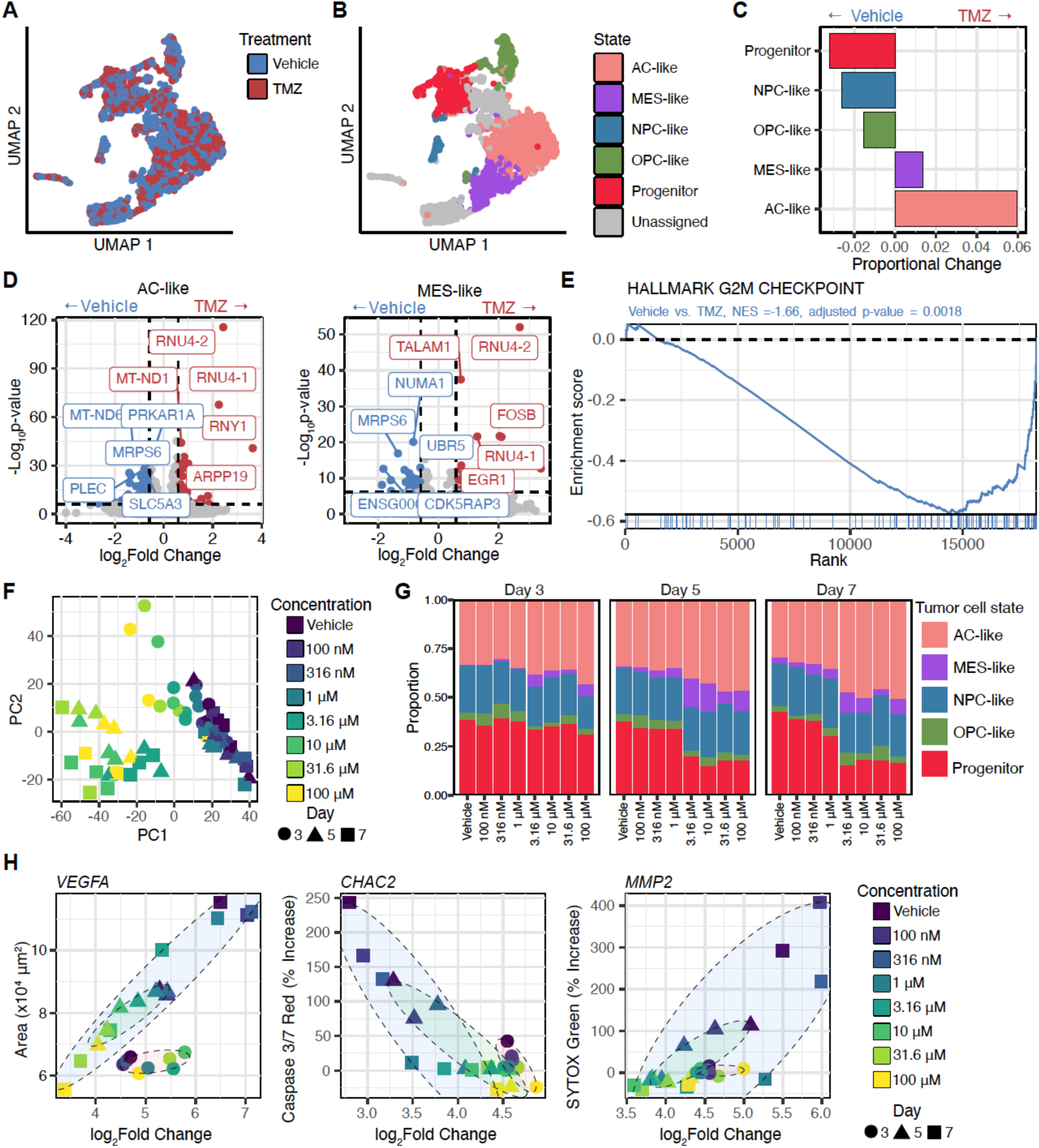
PDN models as a dynamic and heterogeneous platform to explore cellular responses to temozolomide treatment. **A.** Representative UMAP plots of resultant scRNASeq data, colored by treatment type. **B.** Representative UMAP plots of resultant scRNASeq data, colored by annotated cellular state. **C.** Bar plot depicting proportional changes in annotated cellular states between vehicle and temozolomide treated samples. Shifts to the left indicate increases in vehicle populations, whilst shifts to the right indicate increases in temozolomide treated populations. **D.** Volcano plots of DEGs in tumor states increased within temozolomide treated samples. Left, identified DEGs within the AC-like tumor population. Right, identified DEGs within the MES-like tumor population. **E.** GSEA of the hallmark G2M MSigDB gene set performed on the ranked DEG list within the AC-like population as depicted in d), demonstrating negative enrichment within temozolomide treated AC-like cells. **F.** PCA plot of the first and second dimensions of pooled TIRE-Seq samples, demonstrating clear effects across concentration and time. Shape represents day, with color representing concentration. **G.** Bar plot showing deconvoluted tumor states as determined via CYBERSORTx over both dose and time, showing distinct changes in proportions of tumor populations over both time and dose. **H.** Scatter plots of results obtained via LASSO regression for identification of genes whose change in expression correlate with physical PDN measures as quantified by the IncuCyte. Measures for area, Caspase 3/7 Red, and SYTOX green shown.

**Table 2.** Pharmacological characteristics of PDN lines in response to temozolomide, treated from 10 nM – 100 μM in a 9-point dose curve.

| ID | Doubling Time (days) | MGMT Methylation Status | Temozolomide Response (μM) | |
| --- | --- | --- | --- | --- |
|  |  |  | GR<sub>50</sub> | IC<sub>50</sub> |
| GL0016 | 4.13 | Unmethylated | 41.71 μM | 80.44 μM |
| GL0051 | 6.75 | Unmethylated | 77.34 μM | 81.7 μM |
| GL0095 | 5.69 | Methylated | 56.35 μM | 87.55 μM |
| GL0157 | 5.10 | Methylated | 7.22 μM | 9.49 μM |
| BT145 | 2.44 | Methylated | 12.32 μM | 13.07 μM |
| BT504 | 3.24 | Methylated | 2.94 μM | 2.52 μM |

To determine the impact of TMZ longitudinally on tumor cell states, we deconvoluted the cell proportions from the samples over time. Notably, at concentrations above 1 μM and from 5 days drug exposure, AC-like and MES-like tumor cell composition started to increase, at the expense of the progenitor population (**Figure 4G**). This validated the altered cell state in multiple PDN lines and identified the impact of dose and time on the transcriptional signatures of heterogeneous tumor cell states. Due to the bulk nature of TIRE-seq, we were unable to detect individual genes that may be playing a role in this plasticity in each cell state. Thus, future studies examining resistance mechanisms to TMZ should focus specifically on the AC-like and MES-like tumor cell states in patient samples to truly interrogate the impact of therapy in glioma.

## Discussion

Models that reflect patient tumor composition are critical in the drug development pipeline and form the basis of translational research to ultimately improve patient survival. As the understanding of tumor cell state heterogeneity in GBM has emerged, a major question in the field remains selecting an appropriate *in vitro* model to best recapitulate the vast array of cell composition. Here, we leveraged the development of matched cohorts to reveal that both PDO and PDN patient-derived models broadly recapitulate primary tissue samples across genomic, epigenetic and transcriptomic benchmarks. Both models retain tumor cell state heterogeneity consistent with the primary tumor, ensuring pre-clinical utility for drug testing. Validating patient findings, we demonstrate that TMZ impacts tumor cell states in PDN models differently, highlighting the importance of understanding patient heterogeneity in treatment resistance mechanisms.

It is well-established that 2D GBM models are composed primarily of MES-like tumor cells^13,52^ and extended culture conditions can increase *MYC* amplification events^8^, resulting in lines that do not reflect the complex intra-tumoral heterogeneity of primary tissue, hampering pre-clinical studies. Through examination of matched 3D PDO and PDN models, we determined that the key difference with primary tissue is epigenomic and transcriptomic alterations consistent with increased cell proliferation. Critically, tumor heterogeneity, genomic and *MGMT* promoter methylation events remained stable in 3D culture, confirming that either model provides an accurate pre-clinical representation of the patient sample, positioning 3D models as superior platforms for translational research.

Our finding that PDOs, but not PDNs, can contain non-malignant cells, confirms previous studies^14,43^. Surprisingly, we did identify PDOs absent of non-malignant cells, such as in

GL0028. Given the matched primary tumor contained the lowest proportion of non-malignant cells, this suggests that the tumor tissue used to generate the PDO line may have not contained any non-malignant cells as input. Thus, initial sample selection is essential in achieving a mixture of malignant and non-malignant cells in PDO models. Multiple components of the GBM microenvironment have been reported, which include immune, vasculature, extracellular matrix, and in recent times, neurons^14,43^. The bi-directional interactions of these compartments with GBM and associated functional consequences are critical modulators of disease pathogenesis. In a recent co-culture experiment, *Mangena, et al.* ^18^ demonstrated that integration of PDNs within cortical organoids resulted in higher confidence in assignment of cellular states, though this did not alter the overall proportion of states. As such, the presence or absence of non-malignant compartments may potentiate therapeutic responses, a critical consideration when selecting an appropriate model. Notably, we observe that within our cohort, PDOs are established more rapidly compared to PDNs, which may contribute to the preservation of non-malignant cells during the initial stages of model development. The faster establishment of PDOs could potentially provide a more stable microenvironment that supports the maintenance of both malignant and non-malignant cell populations. This confirms previous findings that the presence of non-malignant cells within PDO models can diverge and dilute over time^14,43^. This phenomenon is postulated to be due to culture medium optimized for tumor division, rather than immune expansion.

Patient studies investigating tumor cell states following TMZ treatment have identified increased proportions of MES-like tumor cells, suggesting that this population may be resistant to treatment. To study this phenomenon, pre-clinical models are required that can maintain molecular and cellular heterogeneity over extended periods. We observed increased AC-like states across time and TMZ dose, a finding reproduced in a separate model exposed to a single high dose of TMZ. This approach also identified a lack of change in the inferred proportion of NPC-like cells, an intriguing finding given previous work describes NPC cells as TMZ sensitive^53^. A comprehensive bulk RNA-seq experiment with multiple samples provides more statistical power, broader biological insights, and potentially lower per-sample costs compared to a single-cell RNA-seq approach, which offers high-resolution data on cellular heterogeneity but comes with significantly higher per-sample expenses and more complex computational analysis requirements.

Our analysis reveals that the main source of variation between patient-derived organoid and neurosphere models stems from differences between patient tumors, rather than from the models themselves. This is particularly evident in the distinct clustering of samples in the methylation data, which confirms that the observed heterogeneity is intrinsic to the tumor and not due to the model. Despite some perceived differences, our findings show that both PDOs and PDNs closely resemble the characteristics of the patient tumor. Our data confirms that both patient-derived models recapitulate patient intra-tumoral heterogeneity, providing a platform for tumor cell state refined therapeutic studies.

## Authorship

Z.M., J.R.W., S.F. and S.A.B. conceived the study. O.M.S. and C.S. aided in experimental designs for PDOs. A.F. and K.D. were involved in tissue procurement. Z.M. generated PDN models and C.S. generated PDOs. S.W. and K.L. optimized PDN generation and maintenance protocols. D.B. performed TIRE-Seq experiments. Z.M., A.V. and J.P. maintained PDN cultures. Z.M. and A.V. optimized and extracted nuclei. S.O. and J.P. performed IHC experiments. S.F., S.A.B. and J.R.W. obtained funding and provided supervision. Z.M. undertook all analyses. Z.M., S.F., S.A.B. and J.R.W. wrote and revised the manuscript with input from all authors.

## Data Availability

Data will be made available upon reasonable request.

## Acknowledgements

We thank A. Fakhri, S. Stylli and K. Drummond for expert curation of the Royal Melbourne Hospital Neurosurgery Brain and Spine Tumour Tissue Bank, and D. Zalcenstein and C. Anttila at the WEHI Advanced Genomics Facility.

