## Supplementary Figures 1-4 for "Three-dimensional patient-derived models of glioblastoma retain intra-tumoral heterogeneity"

Supplementary Figure 1

Supplementary Figure 2

Supplementary Figure 3

Supplementary Figure 4

### SUPPLEMENTARY FIGURE 1

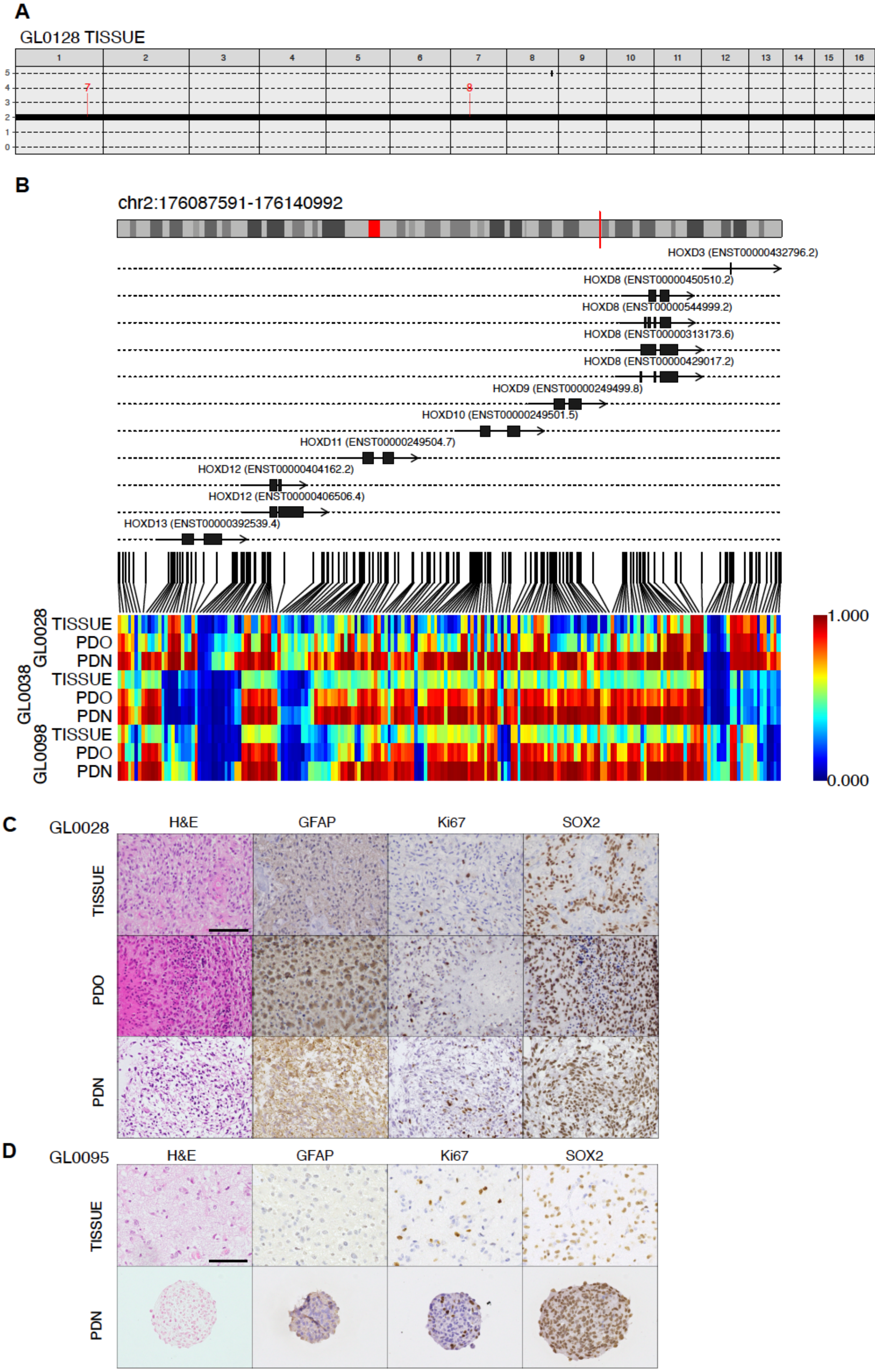

**Supplementary Figure 1 – related to Figure 1**

**A.** Original genomic reference profile for GL0128 tissue. **B.** Differential methylated regions within selected *HOX* genes identified for both PDN and PDO samples**. C.** Representative IHC for GL0028 samples. IHC includes H&E stains, GFAP, Ki67, and SOX2 for PDN, PDO, and tissue samples. Scale bar, 100 μm. **D.** Representative IHC for GL0095 samples. IHC includes H&E stains, GFAP, Ki67, and SOX2 for PDN and tissue samples. Scale bar, 100 μm.

### SUPPLEMENTARY FIGURE 2

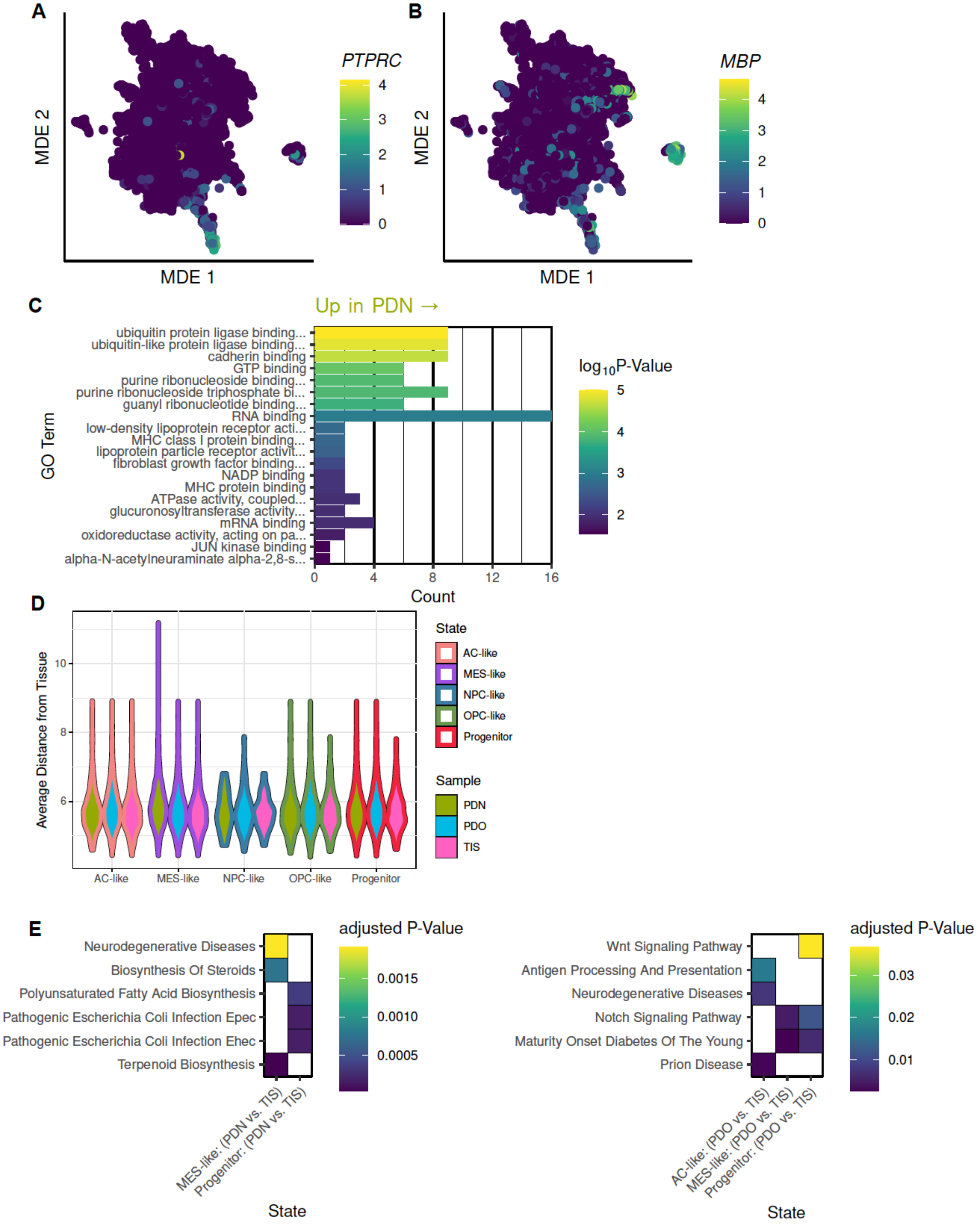

**Supplementary Figure 2 – related to Figures 2 and 3**

**A.** Expression of canonical immune cell marker *PTPRC* across all nuclei. **B.** Expression of canonical oligodendrocyte cell marker *MBP* across all nuclei. **C.** Significantly upregulated GO terms between malignant PDN nuclei and malignant tissue nuclei. **D.** Quantification of average distance between each individual nuclei and all tissue samples within each individual tumor state. Distances were obtained from the network used to create the MDE for all malignant nuclei. **E.** Significantly enriched KEGG pathways for both PDN vs. tissue and PDO vs. tissue comparisons across individual tumor states.

SUPPLEMENTARY FIGURE 3

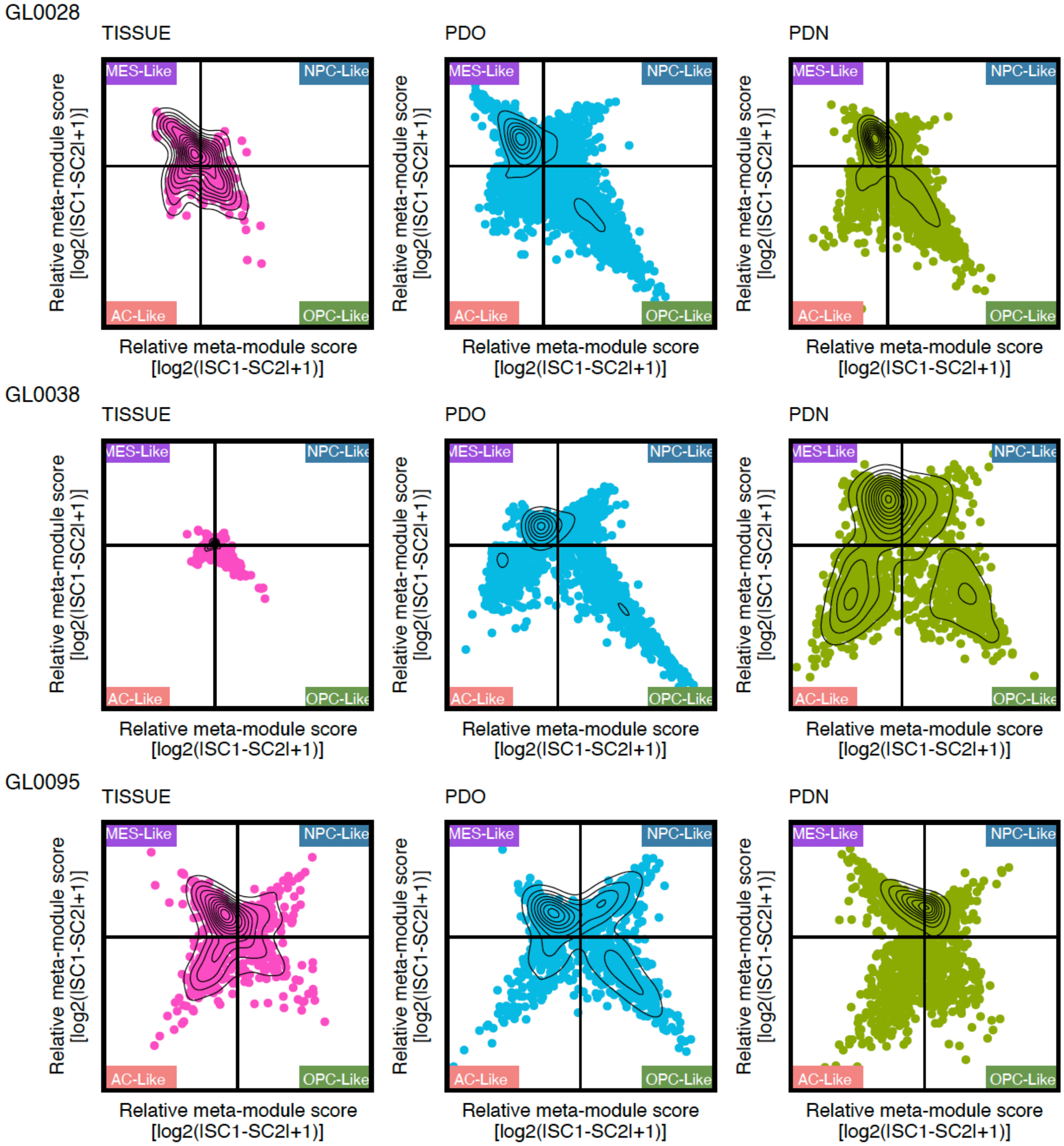


**Supplementary Figure 3 – related to Figure 3**

Meta-module plots of cellular states as defined in *Neftel, et al.* for all samples.

### SUPPLEMENTARY FIGURE 4

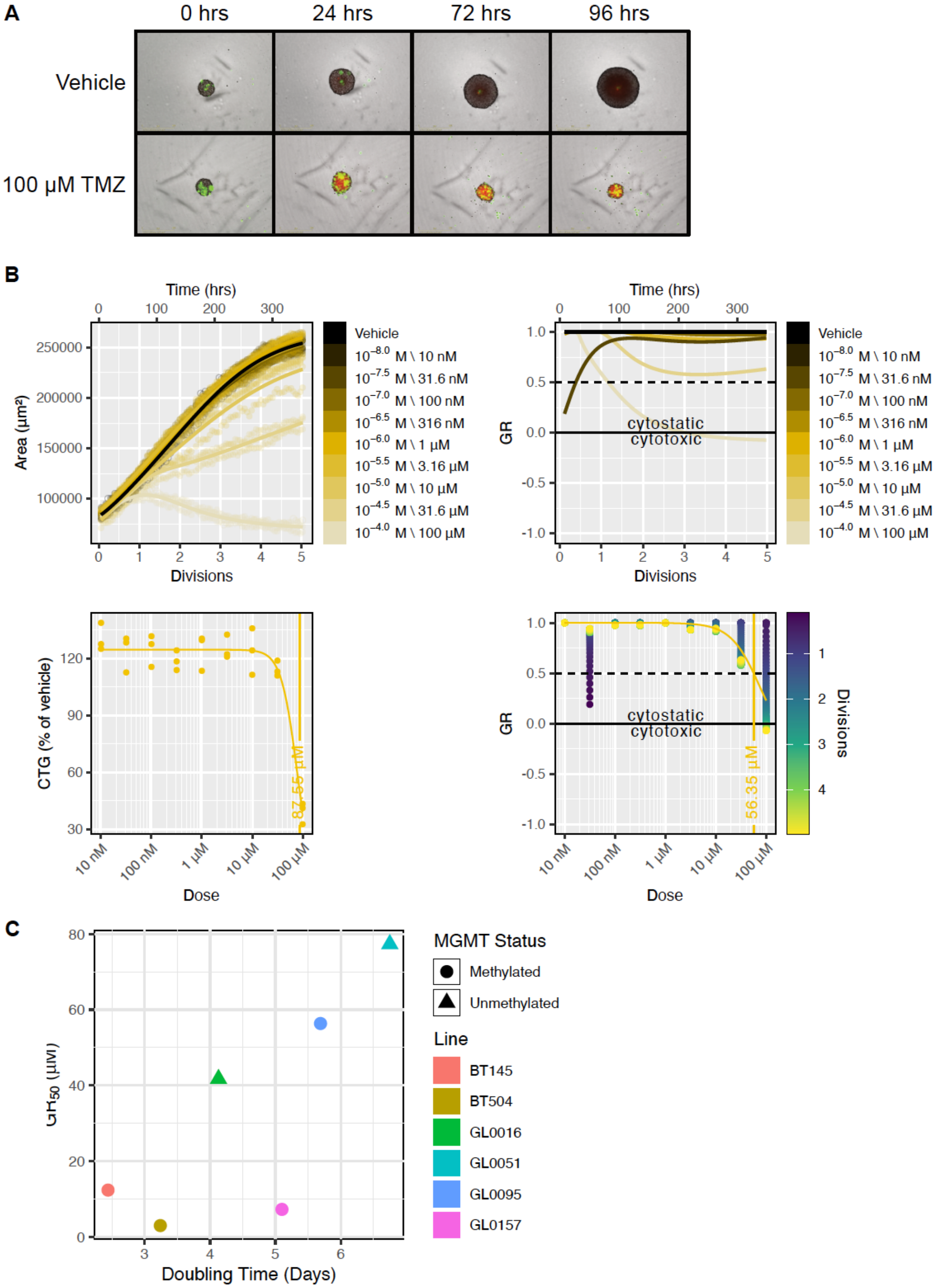

**Supplementary Figure 4 – related to Figure 4**

**A.** Representative IncuCyte images of TMZ treated GL0095 PDN over time. Images are shown as composite, with brightfield images merged with SYTOX Green in green, and Caspase 3/7 in red. **B.** Representative readouts from IncuCyte experiment as shown in (A). In order: final CellTiter-Glo measurement with superimposed $IC_{50}$, segmented area of individual PDN spheres over time, calculated GR values across concentration over time, calculated $GR_{50}$ over time. **C.** Plot demonstrating a lack of correlation between PDN doubling time and calculated $GR_{50}$ values as determined via live-cell imaging. Colors represent individual PDN lines, whilst shapes represent MGMT status.
